# Conservation of EMT transcription factor function in controlling pluripotent adult stem cell migration *in vivo* in planarians

**DOI:** 10.1101/080853

**Authors:** Prasad Abnave, Ellen Aboukhatwa, Nobuyoshi Kosaka, James Thompson, Mark A. Hill, A. Aziz Aboobaker

## Abstract

Migration of stem cells underpins the physiology of metazoan animals. For tissues to be maintained, stem cells and their progeny must migrate and differentiate in the correct positions. This need is even more acute after tissue damage by wounding or pathogenic infections. Inappropriate migration also underpins the formation of metastasis. Despite this, few mechanistic studies address stem cell migration during repair or homeostasis in adult tissues. Here, we present a shielded X-ray irradiation assay that allows us to follow stem cell migration in the planarians. We demonstrate that we can use this system to study the molecular control of stem cell migration and show that *snail* and *zeb-1* EMT transcription factors homologs are necessary for cell migration to wound sites and for the establishment of migratory cell morphology. Our work establishes planarians as a suitable model for further in depth study of the processes controlling stem cell migration in vivo.

## INTRODUCTION

Regeneration and tissue homeostasis in multicellular animals are a result of the activity of their stem cells. Most animal adult life histories include some potential to regenerate lost cells, tissues and organs but the efficiency and extent of the regenerative process varies greatly amongst species. Many basal invertebrates like cnidarians, flatworms and annelids are capable of whole body regeneration and some of these are now experimentally tractable model organisms for studying regeneration and homeostasis (Galliot, 2012; Gehrke and Srivastava, 2016; Tanaka and Reddien, 2011). Studies of the invertebrate stem cells that contribute to regeneration and homeostasis inform us about the origins of key stem cell properties. These include potency, self-renewal, production of the correct quantity and type of progeny, and the interpretation of positional information to ensure regenerated tissue is patterned and functionally integrated. So far few studies in regenerative models have investigated cell migration in vivo in adult animals, even though migration to sites of injury or homeostatic activity is a key stem cell activity for regeneration and repair, and has important biomedical applications (Bradshaw et al., 2015; Guedelhoefer and Sánchez Alvarado, 2012a; Reig et al., 2014). The over-activity of migratory mechanisms is a feature of tumor tissue invasion and the pathology caused by cancers (Friedl and Gilmour, 2009; Friedl et al., 2012). Defects in stem cell migration are likely to contribute to many age-related processes leading to disease. These links remain poorly described, particularly *in vivo* (Goichberg, 2016). Many studies have revealed common mechanisms that drive cell migration in different contexts (Friedl and Alexander, 2011; Friedl et al., 2012; Goichberg, 2016; Ridley et al., 2003). However, studying cell migration *in vivo* is technically challenging, and simple model systems amenable to functional study may have a lot to offer. For example, *in vivo* studies in both *Drosophila* and *C. elegans* during embryogenesis and larval development have proven very useful for unveiling fundamental molecular mechanisms also used by vertebrates (Geisbrecht and Montell, 2002; Hagedorn et al., 2013; Montell, 2003; Reig et al., 2014; Sato et al., 2015). The planarian system, in which pluripotent adult stem cells (known as neoblasts, NBs) and their progeny can be easily identified and studied, is another potentially tractable system for studying cell migration (Eisenhoffer et al., 2008). In particular, planarians offer the potential to study stem cell migration in both an adult and highly regenerative context.

Here we have used the model planarian *Schmidtea mediterranea* to establish methods to study cell migration and show that NB and progeny migration utilize epithelialmesenchymal transition (EMT) related mechanisms in response to tissue damage. To date relatively little focus has been given to stem cell migration in planarians (Guedelhoefer and Sánchez Alvarado, 2012a; Saló and Baguñà, 1985), although it is a necessary component of a successful regenerative outcome. We perfect an assay to allow observation of cell migration and describe several novel phenomena in the planarian system, including homeostatic cell migration mechanisms in the absence of wounding. Migrating cells form extended processes, the frequency of which correlate with cell movement towards the wound site. Using markers of the well characterized epidermal lineage we uncover a close relationship between known NB and progeny lineages and the order and extent of cell migration, demonstrating that cells at some stages of differentiation are more migratory than others. RNAi can be efficiently employed within our migration assay and we demonstrate the requirement for a planarian matrix-metalloprotease, *Smed-MMPa* (*mmpa*), and an ortholog of beta-integrin, *Smed-*β*1-integrin (*β*1-integrin),* for normal cell migration and the formation of extended processes as proof of principle of this approach (Bonar and Petersen, 2017; Isolani et al., 2013; Seebeck et al., 2017). Using RNAi we also show the polarity determinant *Smed-notum* (*notum*) is necessary for homeostatic anterior migration of cells in unwounded animals, but not for cells to form processes or to migrate in response to wounding (Petersen and Reddien, 2011). Our observations of migratory cell behavior and morphology led us to consider if EMT related mechanisms are likely to control cell migration in planarians. We investigated three planarian orthologs of EMT-transcription factors (EMT-TFs) and found that they were all required for stem cell migration in the context of our assay. Our work establishes the conservation of EMT mechanisms controlling cell migration across the breadth of bilaterians and establishes the use of *S. mediterranea* as a highly effective model system to study in vivo adult stem cell migration in a regenerative context.

## RESULTS

### Establishment of an X-ray shielded irradiation assay for tracking stem cell migration

The sensitivity of planarian regenerative properties to high doses of ionizing radiation was established over a century ago (Bardeen and Baetjer, 1904). Later this was attributed to the fact that NBs were killed by irradiation (Wolff, 1962). Partially exposing planarians to ionizing radiation, through use of a lead shield, was shown to slow down regenerative ability and suggested the possibility that NBs were potentially able to move to exposed regions and restore regenerative ability (Dubois, 1949). Recently established methods for tracking cell migration in planarians have either revisited shielding or involved transplanting tissue with stem cells into lethally irradiated hosts (Guedelhoefer and Sánchez Alvarado, 2012a; Tasaki et al., 2016). These methods clearly show movement of NBs and their progeny. We set out with the goal of adapting the shielding approach to establish a practical assay for studying the molecular control of cell migration. We wished to simultaneously use both smaller animals and larger numbers of animals for irradiation to generate a much smaller shielded region and by performing experiments simultaneously across larger numbers of animals, rather than shielding animals individually.

We perfected a technique in which multiple animals can be uniformly irradiated with X-rays, apart from a thin strip in a predetermined position along their body axis. This is achieved by placing the animals directly above a 0.8 mm strip of lead (6.1 mm thick), to significantly attenuate the X-rays in the region just above the lead to less than 5% of the dose in the rest of the animal (Figure 1A-C, Figure S1A-C).

**Figure 1.**
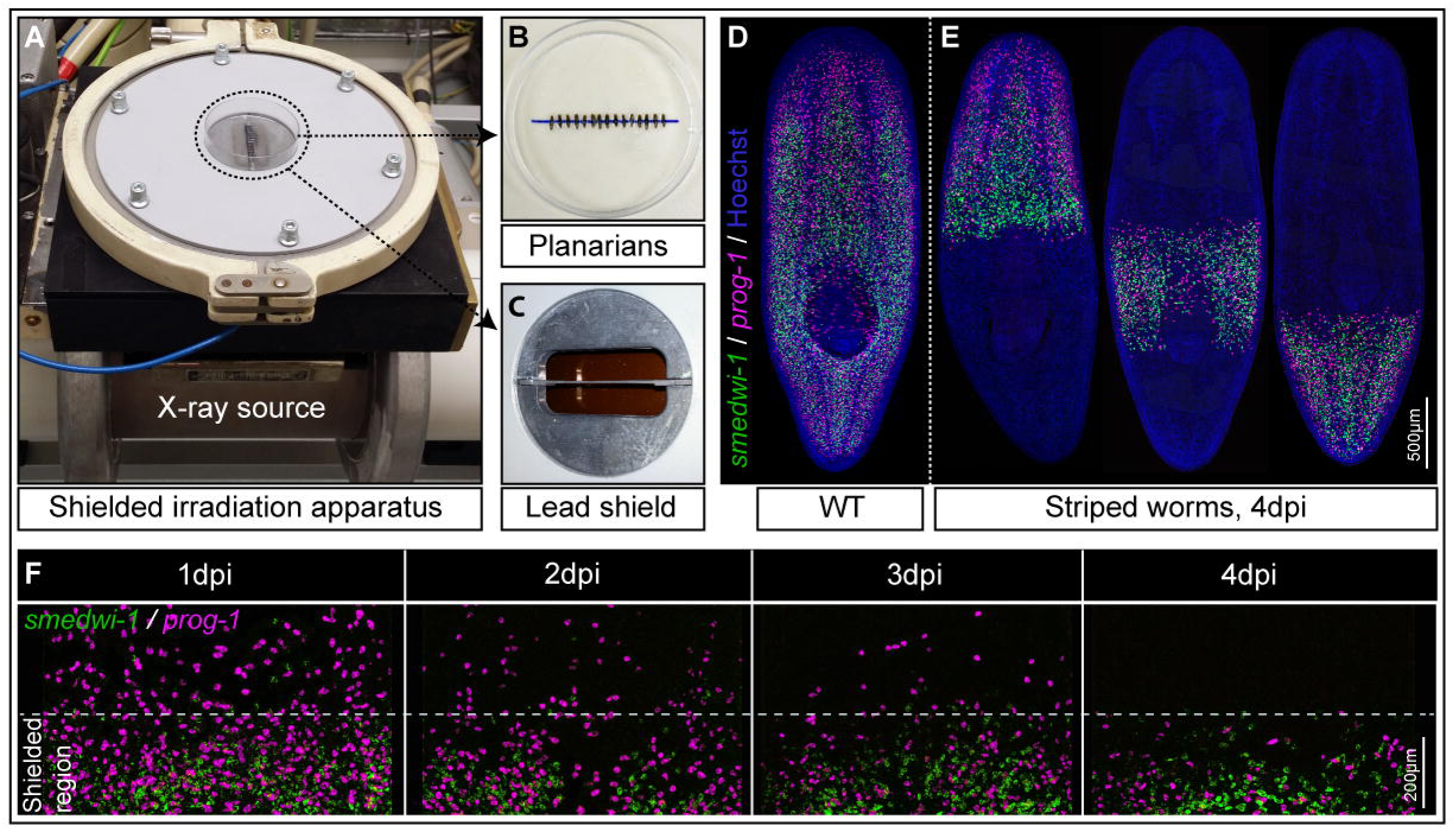
Shielded irradiation assay setup to generate stripped worms. (A) Point source X-ray irradiator with the lead shield on top and holding worms aligned in a Petri dish. (B) Worms anesthetized in 0.2% chloretone and aligned in a straight line on 60mm Petri dish. (C) Lead shield with a horizontal lead stripe in the middle. (D) Wild type un-irradiated planarians showing distribution of NBs (green) and early progeny (magenta). (E) Striped planarians at 4 days post shielded irradiation (4dpi) showing band of stem cells (green) and early progeny (magenta) restricted to the irradiation-protected region. A 30 Gy X-ray does is used. (F) Gradual loss of NBs (green) and early progeny (magenta) in the non-shielded region after 1 dpi, 2dpi, 3dpi and 4dpi respectively (n=10), and maintenance within the shielded region. See also Figure S1.

Our final working version of the apparatus is conveniently designed to fit a standard 60 mm Petri dish, with the lead shield lying below the diameter (Figure 1A, Figure S1A and B). Anaesthetized planarians are aligned across the diameter in preparation for X-ray exposure (Figure 1 A-C). We could then expose up to 20 ∼3-5mm long worms simultaneously to a normally lethal 30 Gy X-ray dose in a 1 min 18 sec exposure, with the shielded region receiving <1.5 Gy. This allows for some precision in controlling the position of a surviving band of NBs (Figure 1D and E).

Looking at animals with the shield positioned centrally along the anterior to posterior (AP) axis we performed whole mount fluorescent in situ hybridization (WFISH) to assay the effectiveness of the shield. With the *smedwi-1* NB marker we confirmed that all NBs (*smedwi-1^+^*) outside the shielded region disappear by 24 hours post irradiation (pi). With the early epidermal lineage marker *prog-1* we confirmed that stem cell progeny (*prog-1^+^*) outside the shielded region have differentiated by 4 days pi and disappear as no NB are present to renew the *prog-1^+^* population (Figure 1E and F). We observed that cells within the shield have a density equivalent to that in wild type animals not subjected to shielded irradiation, suggesting that the shield is effective at protecting cells (Figure 1E and F and see Figure 2D for quantification). We also noted that there is no cell migration from the shielded region during this time (Figure 1E and F). These data established that any observation of migrating NBs and progeny should ideally occur after 4 days pi. In summary, our X-ray shielded assay allows convenient and precise observation of NB and progeny behavior over time post-irradiation, and in animals of a size and number suitable for functional studies.

**Figure 2.**
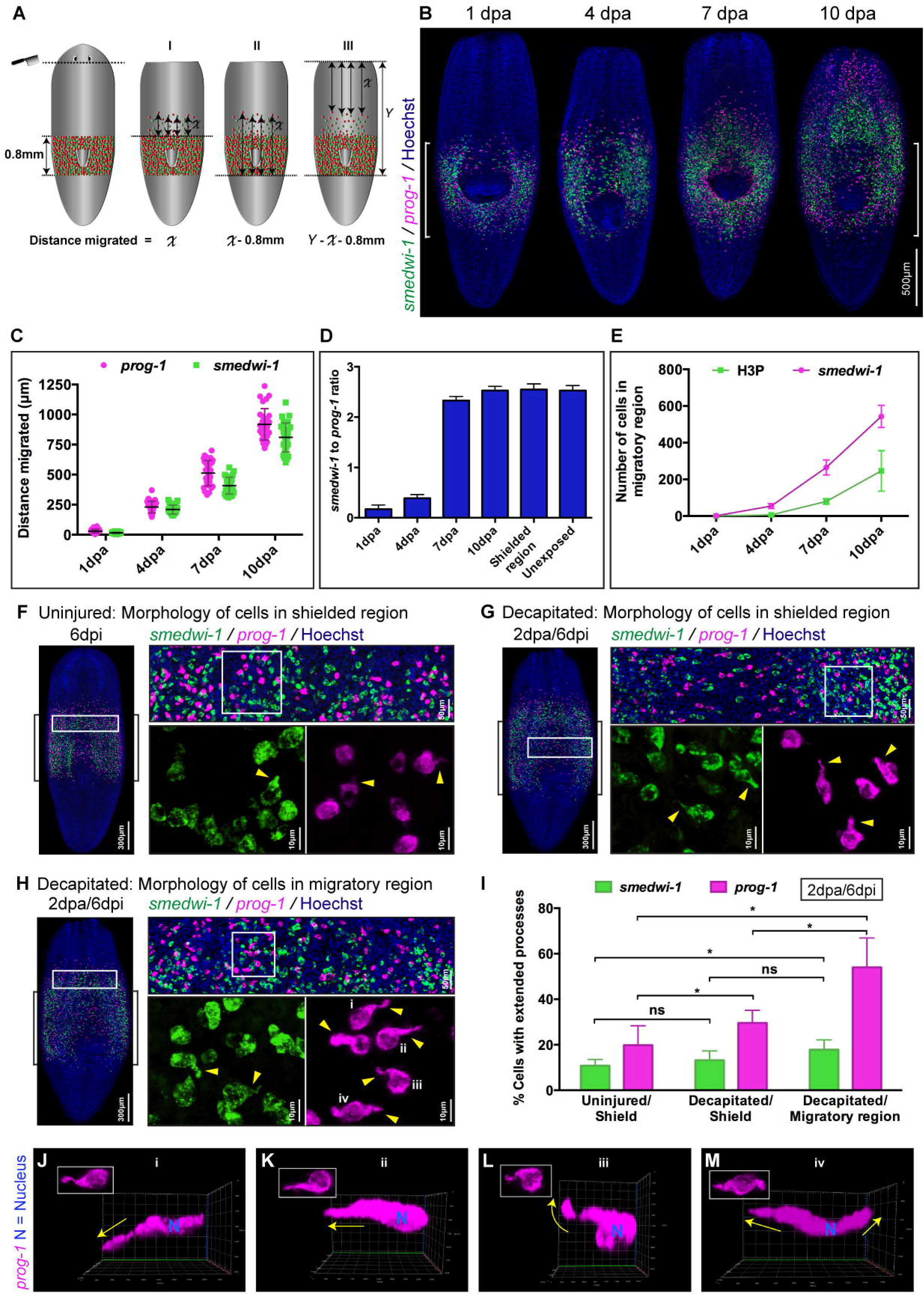
Wound induced cell migration and characteristic extended morphology of migrating stem cells and stem cell progeny. (A) Diagrammatic model demonstrating the position of the wound and three (I, II and III) independent methods for measuring cell migration distances. (B) Representative WFISH showing migration and repopulation of NBs (green) and early progeny (magenta) after shielding of across the pharynx at 1, 4, 7 and 10 days post injury (n=20 per time point). Scale bars: 500μm. (C) Measurements of distances migrated by NBs (green) and early progeny (magenta) at 1, 4, 7 and 10 days post decapitation. Each dot represents average distance migrated by 10 most distal cells in each animal (n=25 per time point). Lines and error bars indicate mean and SD. (D) Numbers of NB to early progeny ratios in the migratory region are plotted at 1, 4, 7 and 10 days post decapitation (n=20 per time point). Ratio of cells in shielded region and in unexposed worms is used as a control. The results are expressed as means ±SD. (E) Quantification of NBs (magenta) and mitotic cells (green) in the migratory region following decapitation at 1, 4, 7 and 10 days (n=20 per time point). The results are expressed as means ±SD. (F) Morphology of cells within the shielded region in an uninjured worm shows very few stem cells (green) and few early progeny cells (magenta) with extended cytoplasmic projections (n=20). (G) Morphology of cells within the shielded region in the decapitated worm shows few stem cells (green) and some early progeny cells (magenta) with extended cytoplasmic projections (n=20). (H) Morphology of cells within the migratory region in the decapitated worm shows few stem cells (green) and many early progeny cells (magenta) with extended cytoplasmic projections (n=20). (I) Quantification shows increase in number of stem cells (green) and early progeny cells (magenta) with extended processes within decapitated/migratory region as well as decapitated/shielded region compared to the uninjured/shielded region (n=20 per condition). The results are expressed as means ±SD. Student’s t test: *p<0.05. (J-M) Early progeny cells (magenta) within migratory region in decapitated worms show extended processes in various directions relative to the wound. Yellow arrows indicate the direction of extended processes. Position of wound relative to cells is to the top. See also Figure S2.

### Features of planarian cell migration after wounding

We next employed the assay system to describe the movement of NBs and progeny. The cycling NBs in *S. mediterranea* are normally present throughout the body but absent from the region in front of the photoreceptors and the centrally positioned pharynx and are not detectable within early regenerative blastema (Figure S2A and B). These facts mean that in normal animals: i) NBs would not normally have far to migrate during normal homeostasis or regeneration as they will always be relatively close to where they are required, except for the anterior region and the pharynx, ii) in the context of early regeneration post-mitotic progeny migrate to establish the blastema tissue before NBs, and iii) at least for the pharynx and the most anterior tissue, homeostasis is achieved by migration of post-mitotic progeny, and not NBs. Together this led us to expect that stem cell progeny might have migratory properties that are distinct from NBs.

We shielded animals over the pharynx (Figure 2A and B) and made anterior wounds by decapitation just under the photoreceptors, at 4 d pi when a ‘blank canvas’ is present anterior to the shielded region (Figure 1F). Using WFISH over a 10 day time course after wounding, we observed that, as previously described, stem cells and stem cell progeny migrated anteriorly towards the wound, but not in a posterior direction (Figure 2B). We used the lack of posterior migration in this experimental design to facilitate accurate measurements of individual cell migration distances over time (Figure 2A). Quantifying *smedwi-1^+^* NBs, *prog1^+^* progeny and mitotic cells in the migratory region just anterior to the pharynx allowed us to develop a detailed overview of the migration process (Figure 2B-E).

While the most advanced *smedwi-1^+^* cells can some times match the extent of migration of the most advanced *prog1^+^* cells, we found that many more *prog1^+^* cells enter the migratory region than *smedwi-1^+^* cells over the first 4 days post amputation (pa). (Figure 2B-D). This observation suggests that progeny react *en masse* to a wound derived signal and NBs follow, either independently in response to the wound signal or because they somehow sense the migration of *prog1*^+^ cells and follow, or some combination of both. By 7 days pa, while the density of NBs and progeny in the migratory region just anterior to the shield are still lower than in unexposed animals, homeostatic ratios of stem cells and stem cell progeny are restored (Figure 2D). We observed cells in M-phase within the field of migrating cells, the numbers of which increased in proportion with the numbers of migrating *smedwi-1^+^* NBs over time (Figure S2C, D and Figure 2E). This pattern of proliferation in the migratory region is consistent with the homeostatic ratio of NBs and progeny being restored by increased stem cell division as well as by further migration from the shielded region (Figure 2C-E). From this we deduce that increases in number of both NBs and progeny outside of the shielded region are fueled initially by migration, but then by both further migration and proliferation of NBs.

*Prog1^+^* progeny that reach the wound site at 10 days pa can only have arisen from asymmetric cell divisions of NBs as old as 6 days pa or later, as 4 days the maximum time before they differentiate and stop expressing the *prog-1* marker (Eisenhoffer et al, 2008). Given the NB migration speeds we observe (Figure 2C), these *prog1^+^* cells must be the progeny of NBs that have themselves already migrated well beyond the shielded region. Taken together, this data suggests that migrating *smedwi-1^+^* NBs undergo both symmetric and asymmetric cell divisions that increase both the number of *smedwi-1^+^* cells and *prog1^+^* cells, importantly providing a source of stem cell progeny that do not derive from the shielded region. We note the overall similarity in these dynamics to that observed during regeneration after amputation, where stem cell progeny form the initial regeneration blastema, with NBs only following later.

We also wished to know how precise the homing of migrating cells to wounds could be. To investigate we performed single poke wounds at the midline or notches confined to one side of the animal (Figure S2E and F). We observed that even these small injuries in relatively close proximity, promoted distinct migratory responses around each wound site, indicating that migrating cells home with precision to injuries (Figure S2E and F). Despite the absence of NBs and progeny in the whole anterior tissue field migrating stem cell progeny only migrate and collect around the wound, and do not sense the absence of NBs and progeny elsewhere (Figure S2E and F). We also observed as a general feature of migration towards the wound site that dorsal *prog1^+^* cells appear to migrate more rapidly than ventral cells to the same wound (Figure S2G and H), and that dorsal *smedwi-1^+^* cells migrate centrally while ventral stem cells migrate across the width of animals (Figure S2I).

### Migrating planarian cells have a distinct morphology of extended cell processes

We next investigated the migrating cells themselves in more detail, to see if we could understand more about how they move in *S. mediterranea*. We imaged migrating cells after wounding and compared them to cells remaining in the shielded region that were static. We observed a significantly higher frequency of individual NBs and progeny with extended cell processes in migratory regions of injured animals than in animals that were uninjured or for cells in the shielded region that were not actively migrating (Figure 2F-I, see Figure S2J and K for different cell morphology). We did not observe any connection or alignment between cells with extended processes, and individual cells migrate independently with rather than any mechanism involving collective cell movement requiring cell-cell junction contact (Friedl and Alexander, 2011; Friedl et al., 2012). This observation suggests that cell migration may involve cellular mechanisms similar to those used during classical EMT (Kalluri and Weinberg, 2009; Lamouille et al., 2014). While net movement of cells is towards the wound site, we note that cell processes can extend in all directions, not just towards the wound (Figure 2J-M). Taken together these data indicate that NBs and progeny respond to wounds with directional precision and by extending cell processes.

### The order and extent of cell migration recapitulates cell lineage

Details of planarian NB and progeny lineages, in particular the epidermal lineage allows detailed tracking of differentiation fates (Eisenhoffer et al., 2008; Tu et al., 2015; van Wolfswinkel et al., 2014). Thus, we can use the cell type markers from these studies to label different populations of NBs and progeny (Figure 3A). We investigated the expression of these markers in migrating cells using a series of overlapping double WFISH experiments. These allowed us to measure the extent of migration of each of these cell types and to observe the relationship between migration and differentiation (Figure 3B-M). We observed that migration distance increases for cells expressing later markers of the epidermal lineage, in particular we see a significant difference in extent of migration between *smedwi-1^+ve^* zeta^+ve^ NBs and *smedwi-1^-ve^* zeta^+ve^ progeny (Figure 3H, I and L). These data suggest that very early post-mitotic progeny may have the highest migratory potential in the epidermal cell lineage. Again, we note that this pattern of differentiation and migration recapitulates early regeneration, where cycling NBs do not enter the blastema, which is first populated by post-mitotic progeny.

**Figure 3.**
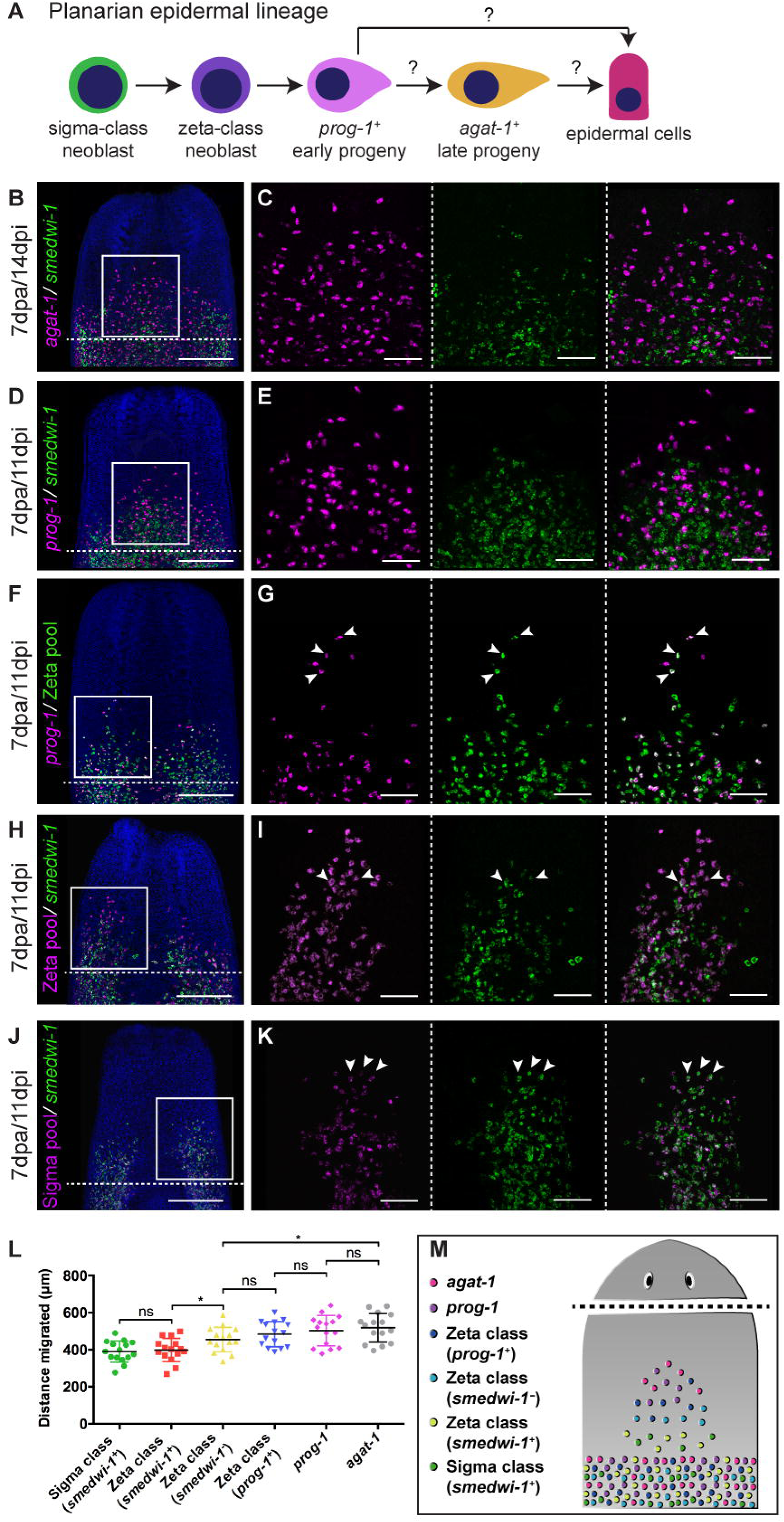
Migration of different epidermal lineage cells shows that cells migrate in the specific order with most differentiated progeny to undifferentiated NBs. (A) Current model of planarian epidermal lineage differentiation. Sigma class NBs give rise to zeta class NBs that produces *prog-1^+^* early progeny. *prog-1^+^* early progeny differentiate into *agat-1^+^* late progeny which terminally differentiate into epidermal cells. (B-K) FISH showing migration of different cell types from epidermal lineage at 7dpa. *agat-1* cells (magenta) migrate way ahead of *smedwi-1* cells (green) (B, C). *prog-1* cells (magenta) migrate way ahead of *smedwi-1* cells (green) (D, E). *prog-1* cells (magenta), Zeta class cells (green) and *prog-1* + Zeta class double positive cells (white) migrate with the similar speed (F, G). *smedwi-1^-^* zeta class cells (magenta) migrate way ahead of *smedwi-1*^+^ zeta stem cells (white) and *smedwi-1* cells (green) (H, I). *smedwi-1*^+^ sigma stem cells (white) and *smedwi-1* cells (green) migrate with the similar speed (J, K) (n=5 per condition). White arrows indicate the examples of double positive cells. Scale bars: 300μm for zoomed out and 100μm for zoomed in view. (L) Measurements of distance travelled by different cell populations (*smedwi-1*^+^ sigma class stem cells, *smedwi-1*^+^ zeta class stem cells, *smedwi-1*^-^ zeta class cells, *prog-1* + zeta class double positive cells, *prog-1* cells and *agat-1* cells) in decapitated worms at 7dpa. (n=15 per condition, Student’s t test: *p<0.05) (M) Model demonstrating order in which different cells migrate following decapitation. *agat-1*, *prog-1*, *prog-1* + zeta class double positive cells migrate most anteriorly and are most distal to the shielded region. Zeta class (*smedwi-1*^-^) cells migrate equally with *prog-1*, *prog-1* + zeta class double positive cells but are quite distant to *agat-1* cells. Zeta stem cells and sigma stem cells migrate with the slowest speed and are most proximal to the shielded region.

### A matrix metalloprotease and beta-integrin are both required for cell migration to wound sites

Having provided a detailed description of cell migration in *S. mediterranea* we next wished to test if we could study gene function in the context of migration. For this we considered candidate genes that might be required for cell migration based on both previous work in planarians and by analogy with other studies of cell migration. This led us to select *mmpa* and β*1-integrin* as strong candidates for proof of principle experiments.

Previous research had attempted to implicate *mmpa*, one of four matrix metalloprotease enzymes identifiable in the *S. mediterranea* genome, as having a possible role in cell migration (Isolani et al., 2013). We decided to look at the function of this gene in the context of our migration assay. We first performed RNAi in the context of normal regeneration and amputation, and observed that *mmpa(RNAi)* animals showed regeneration defects as previously described, with failure to correctly regenerate anterior or posterior tissues (Figure S3A). We then performed RNAi and amputation in the context of our assay and observed that anterior tissues regressed and that animals failed to regenerate (Figure S3B). We used WFISH to monitor the movement of *smedwi-1^+^* NBs and *prog1^+^* stem cell progeny after *mmpa(RNAi)*, and observed almost no migration of cells compared to control *gfp(RNAi)* worms (Figure 4A, D and M, see also Figure S3M and N). Additionally, we examined the morphology of NBs and progeny and observed reduced numbers of cells with extended processes compared to migrating cells in the *gfp(RNAi)* control animals (Figure 4 B, C, E, F and N). These results confirm that this matrix metalloprotease enzyme is required to facilitate cell migration in planarians and demonstrates the potential utility of our assay in generating insights into how stem cell migration is controlled. We found that *mmpa* is only expressed at relatively low levels in stem cells and stem cell progeny, with the bulk of its expression in differentiated radiation insensitive cells (Figure S3C-E) (Kao et al., 2017). We also did not detect *mmpa* expression in migrating cells (Figure S3F and G), suggesting it is instead produced by differentiated cells and required in the extracellular matrix to allow cell extensions to form and allow migration.

**Figure 4.**
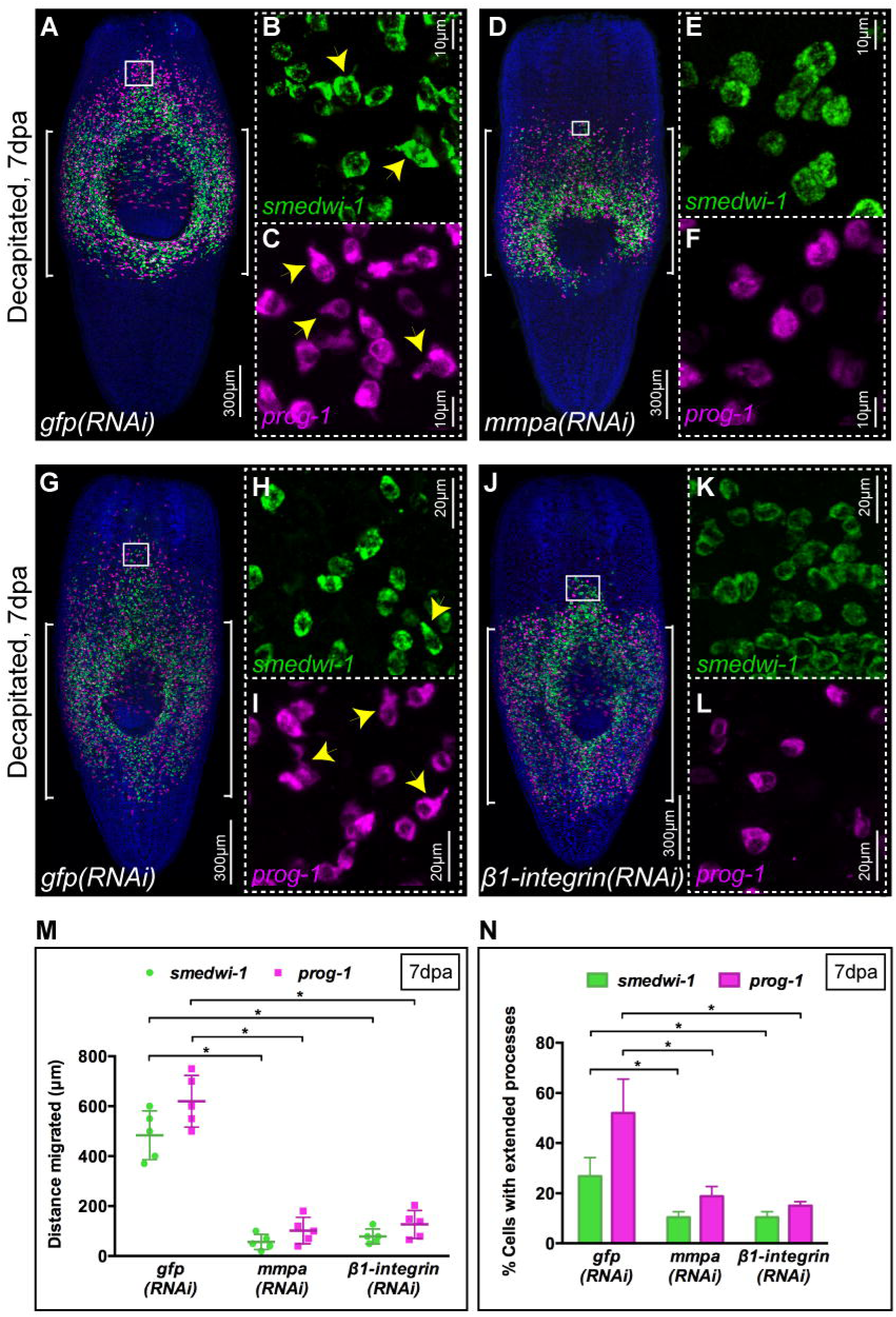
*mmpa* and β*1-integrin* are essential for stem cell and progeny migration as well as for developing extended cell morphology. (A-L) FISH shows migration of stem cells (green) and early progeny (magenta) at 7dpa in *gfp(RNAi)* (A-C, G-I) worms but the migration is inhibited in *mmpa(RNAi)* (D-F) as well as in β*1-integrin(RNAi)* (J-L) worms. Insets show the presence of stem cells (green) and early progeny (magenta) with extended cytoplasmic projections in migratory region of *gfp(RNAi)* worms (B, C, H, I) but are almost absent in *mmpa(RNAi)* (E, F) and β*1-integrin(RNAi)* (K, L) worms (n=5). (M) Measurements shows drastic decrease in the distance migrated by stem cells (green) and early progeny (magenta) at 7dpa in *mmpa(RNAi)* and β*1-integrin(RNAi)* animals compared to *gfp(RNAi)* worms (n=5). Each dot represents the average distance migrated by 10 most distal cells from each animal. Lines and error bars indicate mean and SD. Student’s t test: *p<0.05. (N) Quantification shows that stem cells (green) and early progeny (magenta) with extended processes are reduced significantly in *mmpa(RNAi)* and β*1-integrin(RNAi)* animals in comparison with *gfp(RNAi)* animals at 7dpa (n=5). The results are expressed as means ±SD. Student’s t test: *p<0.05. See also Figure S3.

We next investigated whether β*1-integrin* also had a conserved role in allowing cell migration in our assay. Integrins have conserved roles in orchestrating cell migration, providing a connection between physical actions of the actin cytoskeleton and signaling mechanisms instructing migratory activity (Mogilner and Keren, 2009; Vicente-Manzanares et al., 2009). A consideration of the recently published regenerative phenotypes for planarian β*1-integrin* suggested to us that the cellular disorganization observed in these studies could be in part due to failures in migratory activity (Bonar and Petersen, 2017; Seebeck et al., 2017). We observed that β*1-integrin* transcript was expressed in nearly all *smedwi-1^+^* NBs and about a third of migrating progeny in the migration region of wildtype animals in our assay (Figure S3H-L). We performed (*β1-integrin)RNAi* and found that cell migration was greatly impaired compared to *gfp(RNAi)* controls (Figure 4G-N, Figure S3M and N). Cell process formation in NBs and progeny was also disrupted (Figure 4K, L and N). These data confirm a conserved role for β*1-integrin* in NB and progeny cell migration in planarians, and along with the *mmpa(RNAi)* phenotype confirm that our assay can be combined with RNAi based loss of function studies.

### Anterior migration of stem cells and stem cell progeny in the absence of wounding

While wounding will trigger migration, and in fact precise homing of NBs and progeny (Figure S2 E and F), we wished to observe what happened in the absence of wounding. We shielded animals of equal size at different positions along the AP axis and irradiated them (Figure 5A). When the shield was placed in the posterior region of worms we observed tissue death and regression from the anterior towards the shield (Figure 5B). Subsequently, we observed blastema formation and normal regeneration that took up to 50 d pi (Figure 5C). Using WFISH we were able to observe that NBs and progeny did not migrate until the regressing anterior tissue boundary was relatively close to the anterior of the shielded region (Figure 5D). When animals where shielded in mid body regions with the top of the shield level with the most anterior region of the pharynx we observed regression of the anterior and posterior tissue (Figure 5E). We subsequently observed blastema formation and regeneration that took up to 45 d pi (Figure 5F). WFISH revealed that in these animals NBs and progeny migrate towards the anterior (Figure 5G) and later towards the posterior once regressing tissue is close to the shielded regions. These data suggested that remaining NBs maintain local tissue homeostasis, and remain stationary within the shielded region until regressing tissue boundaries are close enough to trigger migration.

**Figure 5.**
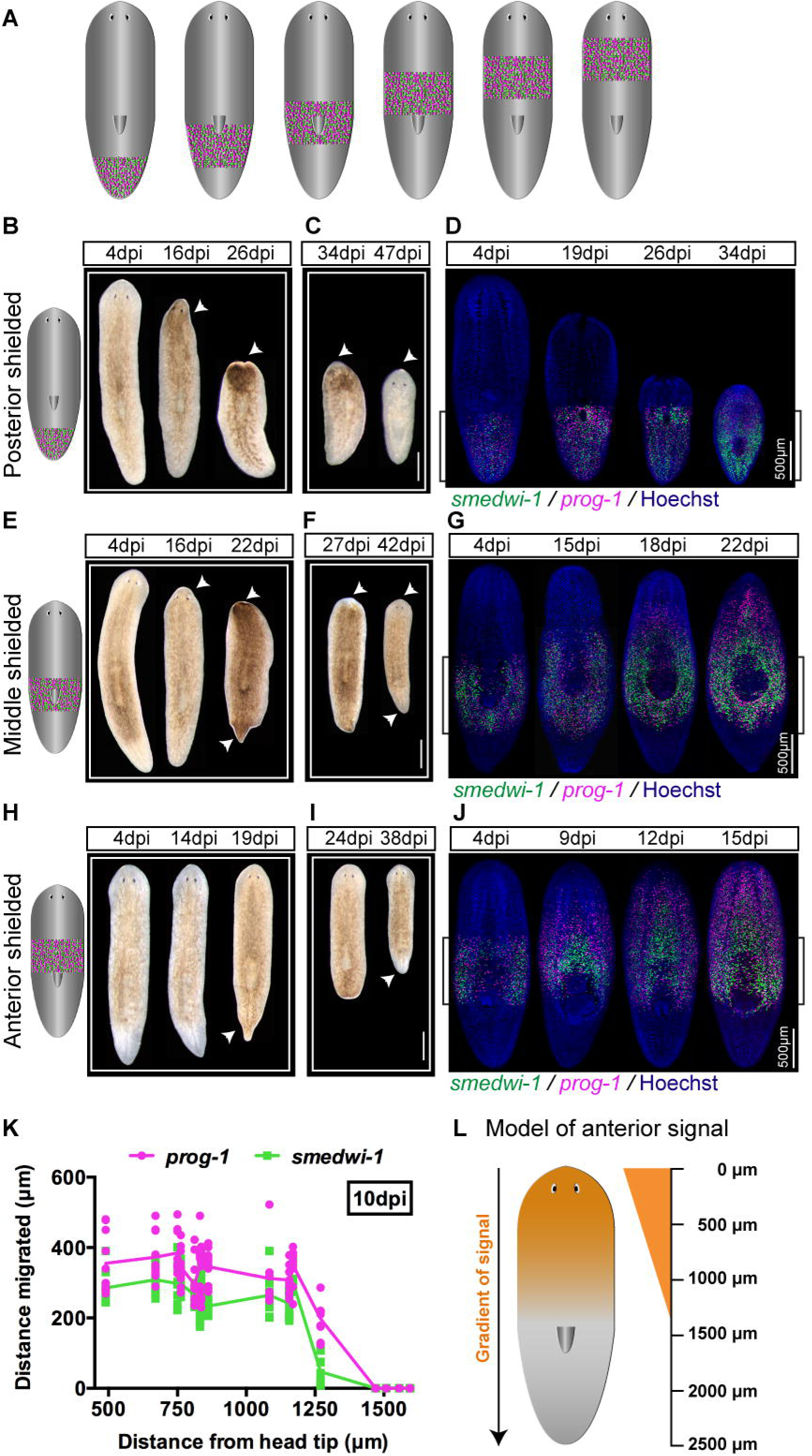
Stem cells and progeny characteristically migrate in anterior direction even without an injury. (A) Cartoon showing strategy of shielding worms at various places along the anterior-posterior axis. (B-J) Bright field images of worms shielded at 3 different places, posterior (B, C), middle (E, F) and anterior (H, I) in shielded irradiation assay showing regression and recovery over the time. Bright field images show head regression in posteriorly shielded worms (B), head and tail regression in middle shielded worms (E) and tail regression in anteriorly shielded worm (H). As cells migrate and repopulate the regressed anterior and posterior regions recovered over the time in all posterior (C), middle (F) and anterior (I) shielded worms (n=20 per time point). Scale bars: 500μm. (D, G, J) FISH showing no migration of stem cells (green) and early progeny (magenta) in posteriorly shielded worms (D) until anterior tissue regress close enough to the shielded region. Whereas, stem cells (green) and early progeny (magenta) migrate as well as repopulate towards the anterior direction in middle (G) and anteriorly (J) shielded worms (n=20 per time point). Migration takes less time in anteriorly placed shields. Scale bars: 500μm. (K) Measurements of distance migrated by stem cells (green) and early progeny (magenta) in the worms shielded irradiated at different places along AP axis. Each dot represent average distance migrated by 10 most distal cells in each animal (n=6). (L) Model showing gradient of signal (orange) form head tip to up to ∼1300μm towards posterior.

In contrast, for animals where the posterior of the shield was positioned level with the anterior of the pharynx we observed that worms often displayed posterior regression but not anterior regression (Figure 5H and I). The heads of these animals never regressed while tails regressed and then regenerated over several weeks (Figure 5I). WFISH subsequently revealed that NBs and progeny could migrate towards the anterior in the absence of wounding or loss of tissue homeostasis (Figure 5J). These results suggest that leaving a stripe of more anteriorly positioned cells is somehow sufficient to trigger anterior migration and maintain anterior tissue homoestasis.

To investigate this phenomenon further we irradiated animals with shields positioned at different points along the AP axis and performed WFISH to observe NBs and stem cell progeny migration at different time points. We were able to observe migration of cells towards the anterior in the absence of wounding as long the shield was within a set distance of the anterior tip (up to 1.2 mm in animals 3 mm in length, Figure 5K and L). These data add to previous work that described that migration only occurs after wounding or when tissue homeostasis fails and tissue regression reaches remaining stem cells (Guedelhoefer and Sánchez Alvarado, 2012a). We find that when stem cells and stem cell progeny in the pre-pharyngeal anterior region can migrate to the anterior in the absence of wounding and before tissue homeostasis fails. This migratory activity restores the normal anterior distribution of both NBs and progeny, suggesting the presence of anterior signals that can call NBs and progeny into the brain and anterior structures over a restricted range. These observations suggest that an anterior signal exists for encouraging cell migration in intact animals that acts at least over the brain region (Figure 5L).

### *Notum* is required for anterior cell migration in intact animals, but not after wounding

By analogy with other systems there are clearly a large number of conserved candidate signaling pathways that could be involved in promoting cell migration. We chose to study two candidate molecules, *Smed-wnt1* (*wnt1*) and *notum* that are both upregulated at anterior wounds in planarians (Petersen and Reddien, 2009). In addition, *notum* is also expressed at the anterior medial tip of intact animals (Petersen and Reddien, 2011) and is therefore also a candidate for controlling anterior migration in the absence of wounding.

It has been previously shown that wounding at any sites results in the transcriptional expression of *wnt1* in muscle cells at the wound site (Witchley et al., 2013). Given that Wnt signaling has a role in regulating cell migration elsewhere (Mayor and Theveneau, 2014), Wnt1 resulting from wound-induced expression could be required for cell migration to the wound in planarians. We performed *wnt1(RNAi)* and observed full penetrance of the tailless phenotype previously described for these animals (Figure S4A) (Petersen and Reddien, 2009). After shielded irradiation we also observed *wnt1(RNAi)* animals were able to regenerate anterior structures completely (Figure S4B). Using WFISH we observed no effects on either NB or progeny migration after wounding, and both cell populations formed cell extensions to a similar extent to control *gfp(RNAi)* animals suggesting that *wnt1(RNAi)* has no essential role in the migration process (Figure 6A-C and G-K).

**Figure 6.**
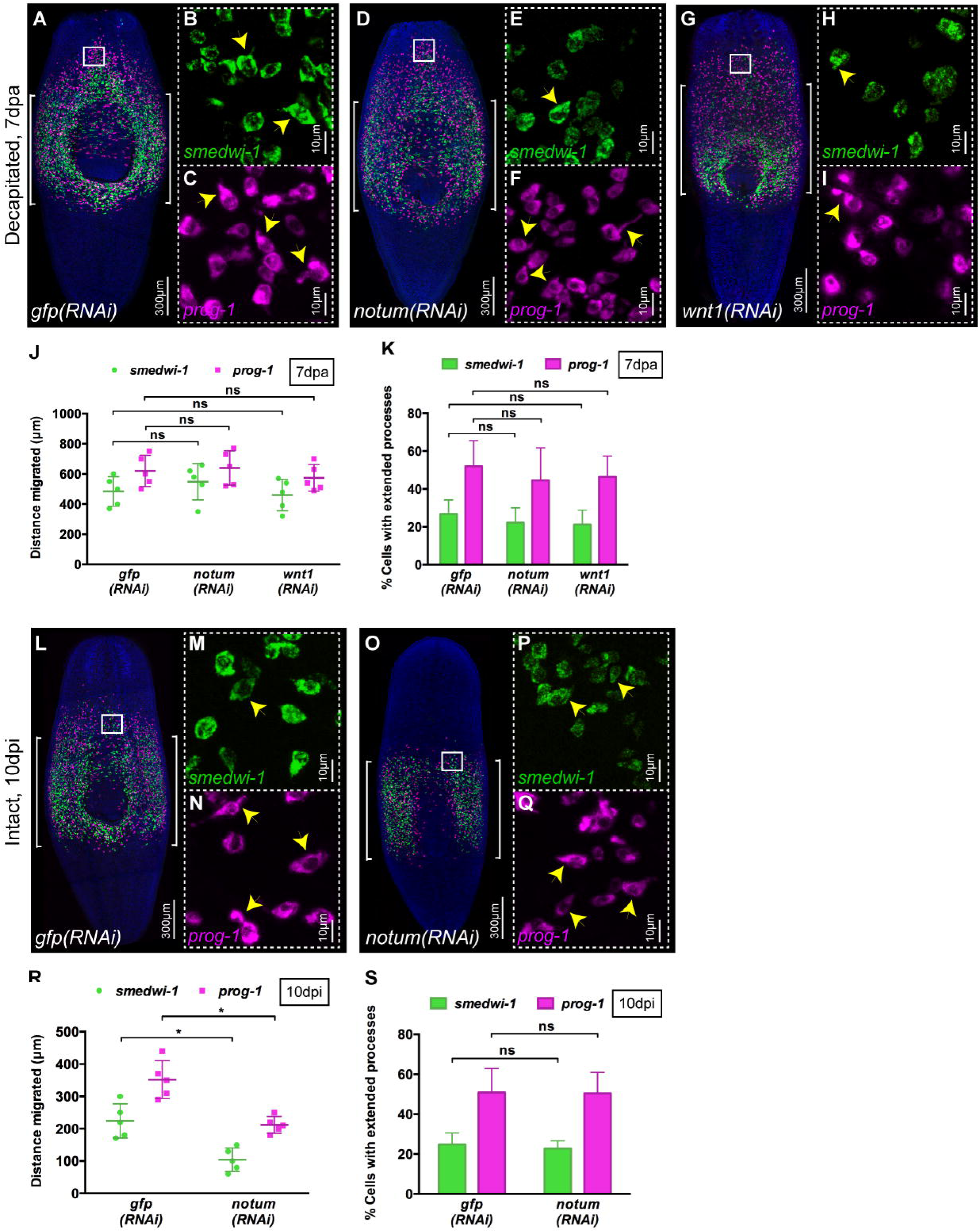
Effect of *notum* RNAi and *wnt1* RNAi on cell migration. (A-I) FISH showing migration of stem cells (green) and early progeny (magenta) at 7 days post decapitation in *gfp(RNAi)* (A-C) animals and is unaffected in *notum(RNAi)* (D-F) and *wnt1(RNAi)* (G-I) animals. Insets shows that stem cells (green) and early progeny (magenta) in migratory region from *gfp(RNAi)* (B, C), *notum(RNAi)* (E-F) and *wnt1(RNAi)* (H-I) are able to form extended processes. (J) Measurements show that distance migrated by stem cells (green) and early progeny (magenta) at 7 dpa in *gfp(RNAi)*, *notum(RNAi)* and *wnt1(RNAi)* animals is equal (n=5). Each dot represents the average distance migrated by 10 most distal cells from each animal. Lines and error bars indicate mean and SD. Student’s t test used for analysis. (K) Quantification shows that the number of stem cells (green) and early progeny (magenta) with extended processes is unaffected in *notum(RNAi)* and *wnt1(RNAi)* animals compared to *gfp(RNAi)* animals (n=5). The results are expressed as means ±SD. Student’s t test used for analysis. (L-Q) FISH showing reduced migration of stem cells (green) and early progeny (magenta) at 10dpi in intact *notum(RNAi)* (O-Q) animals compared to intact *gfp(RNAi)* (L-N) animals. Insets show extended morphology of stem cells (green) and early progeny (magenta) in migratory region (M, N, P, Q). (R) Measurements show that distance migrated by stem cells (green) and early progeny (magenta) at 10dpi in *notum(RNAi)* animals is significantly reduced compared to *gfp(RNAi)* animals (n=5). Each dot represents the average distance migrated by 10 most distal cells from each animal. Lines and error bars indicate mean and SD. Student’s t test: *p<0.05. (S) Quantification shows that the number of stem cells (green) and early progeny (magenta) with extended processes is unaffected in *notum(RNAi)* compared to *gfp(RNAi)* animals (n=5). The results are expressed as means ±SD. Student’s t test used for analysis. See also Figure S4.

*Smed-notum* is also expressed in muscle cells on wounding, but only at anterior facing wounds where it is required to ensure the proper specification of anterior fates, probably by repressing Wnt signaling (Petersen and Reddien, 2011). Additionally it has a homeostatic expression pattern at the anterior margin and has previously been shown to promote the homeostasis and correct size of the brain in combination with the activity of a *wnt11-6* gene expressed in posterior brain regions (Hill and Petersen, 2015). On this basis *notum* represents a candidate molecule for both wound-induced migration and migration of cells towards anterior regions in uninjured animals. We performed *notum(RNAi)* and observed full penetrance of the double tailed phenotype previously described for these animals in a standard regeneration assay (Figure S4A) (Petersen and Reddien, 2011). After shielded irradiation and wounding we observed that while *notum(RNAi)* animals failed to regenerate normal anterior structures compared to controls, we observed no difference in migration of cells or migrating cell morphology compared to control *gfp(RNAi)* animals using WFISH (Figure 6A-F, J and K). However, when using an anteriorly positioned shield, which led to anterior migration of cells in control intact unwounded *gfp(RNAi)* animals, we observed a significant reduction in anterior migration after *notum(RNAi)* (Figure 6L-S, Figure S4C-E). This reduction in migration was not accompanied by a difference in the number of cells with cell extensions (Figure 6S), suggesting that *notum* may act by contributing a directional signal rather than controlling cellular migratory behavior of anteriorly positioned NBs and progeny. These data suggest that *notum* is not essential for wound-induced cell migration but is required in the case of homeostatic anterior migration in intact animals that we uncovered in this work. It seems likely that an earlier description of a *notum/wnt11-6* regulatory circuit involved in homeostatic regulation of brain size may also have a broader role in the homeostatic maintenance of anterior regions that do not normally contain NBs (Hill and Petersen, 2015).

### Conserved EMT transcription factors regulate cell migration in planarians

We next considered if we could establish a broad comparative context for the control of cell migration in planarians and migration in other systems, including mammals. Our observation that NBs and progeny appear to migrate individually using cell extensions to interact with the extracellular matrix and non-migratory neighboring differentiated cells suggested that they may use similar mechanisms to those attributed to EMT (Thiery and Sleeman, 2006). EMT in different contexts requires the activity of a restricted group of transcription factors (EMT-TFs) (Batlle et al., 2000; Cano et al., 2000; Colvin Wanshura et al., 2011; Lamouille et al., 2014). In planarians we identified 2 members of the *snail* transcription factor family (*snail-1* and *snail-2*) of EMT-TFs and an ortholog of the Zinc finger E-box binding homeobox 1 (*zeb-1*) EMT-TF.

We decided to test whether any of these conserved EMT-TF genes were involved in cell migration in planarians. Previously a snail family transcription factor, *snail-2*, has been reported as being expressed in collagen positive muscle cells, in a small percentage of G2/M NBs before wounding and in ∼35% of G2/M NBs after wounding (Scimone et al., 2014).To our knowledge no phenotype has been reported for a snail family gene in planarians and when we performed both *snail-1(RNAi)* or *snail-2(RNAi)* with a standard regenerative assay we observed no phenotypes, and all animals regenerated normally (Figure S5A).

When we performed *snail-1(RNAi)* or *snail-2(RNAi)* in the context of our migration assay, animals failed to regenerate after wounding suggesting a defect in cell migration (Figure S5B). Using WFISH experiments we observed a clear decrease in the extent of cell migration compared to *gfp(RNAi)* animals (Figure 7A, D, G and P, Figure S5M and N). This defect in migration of both NBs and progeny was accompanied by a decrease in the number of cells with cell extensions (Figure 7B, C, E, F, H, I and Q).

**Figure 7.**
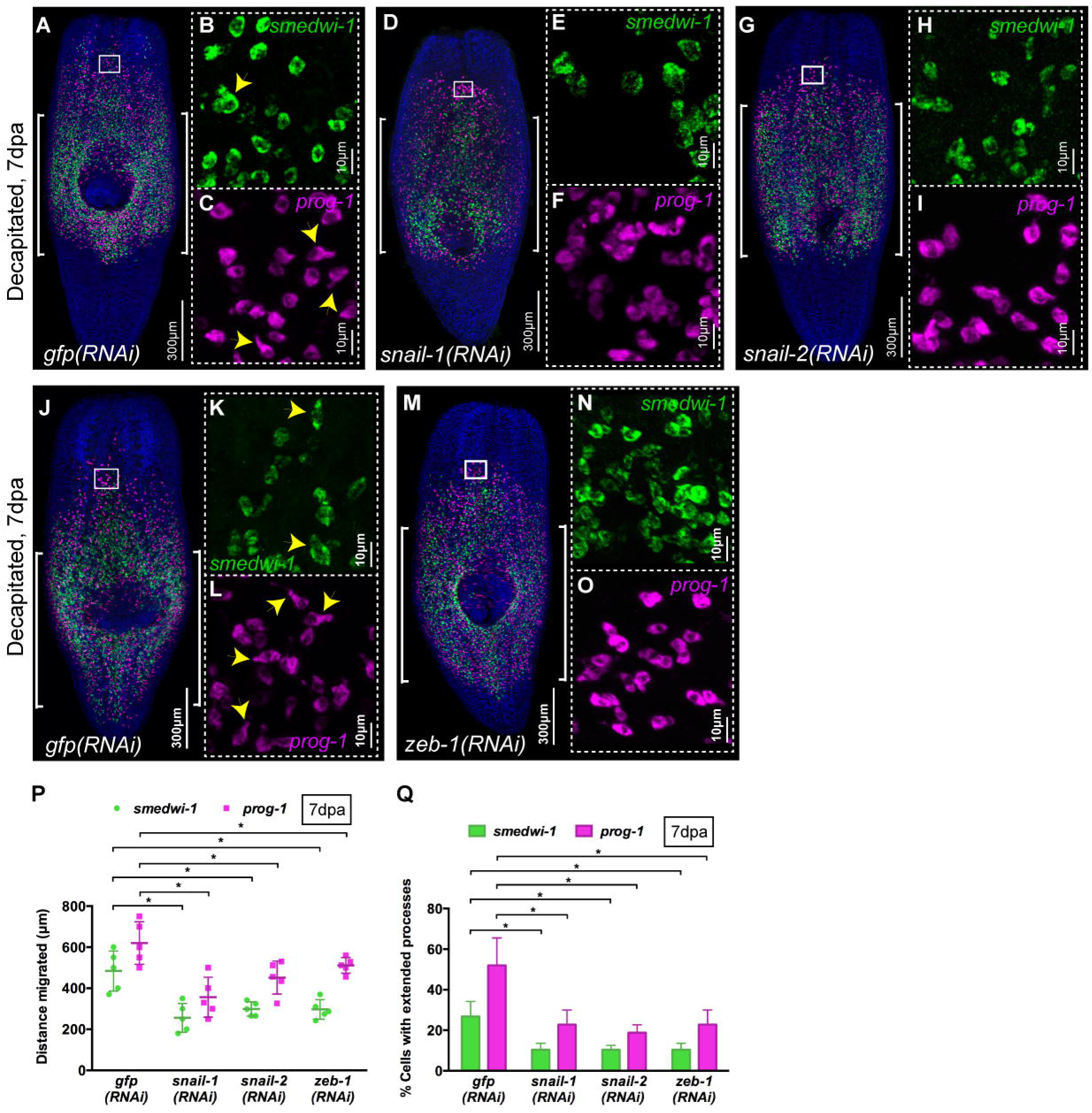
Snail family genes control stem cell and progeny migration. (A-O) FISH shows migration of stem cells (green) and early progeny (magenta) at 7dpa in *gfp(RNAi)* (A-C, J-L) worms but the migration is inhibited in *snail-1(RNAi)* (D-F), *snail-2(RNAi)* (G-I) as well as in *zeb-1(RNAi)* (M-O) worms. Insets show the presence of stem cells (green) and early progeny (magenta) with extended cytoplasmic projections in migratory region of *gfp(RNAi)* worms (B, C, K, L) but are reduced in *snail-1(RNAi)* (E, F), *snail-2(RNAi)* (H-I) and *zeb-1(RNAi)* (N-O) worms (n=5). (P) Measurements shows drastic decrease in the distance migrated by stem cells (green) and early progeny (magenta) at 7dpa in *snail-1(RNAi)*, *snail-2(RNAi)* and *zeb-1(RNAi)* animals compared to *gfp(RNAi)* (n=5). Each dot represents the average distance migrated by 10 most distal cells from each animal. Lines and error bars indicate mean and SD. Student’s t test: *p<0.05. (Q) Quantification shows that stem cells (green) and early progeny (magenta) with extended processes are reduced significantly in *snail-1(RNAi)*, *snail-2(RNAi)* and *zeb-1(RNAi)* animals in comparison with *gfp(RNAi)* at 7dpa (n=5). The results are expressed as means ±SD. Student’s t test: *p<0.05. See also Figure S5 and S6.

We found that both *snail-1* and *snail-2* were expressed in most *smedwi-1^+^* NB cells in the migratory region after wounding (87% and 93% respectively) (Figure S5F and K). This expression patterns suggest that these EMT- have a cell autonomous role in controlling NB migration. Taken together our data suggest that cell autonomous migratory mechanisms are affected by *snail-1(RNAi)* and *snail-2(RNAi)* and establish that *snail* EMT-TFs in planarians have a conserved role in regulating cell migration in response to wound signals.

We also investigated the role of *zeb-1* and similar to our observations for *snail* genes, no defects were observed in *zeb-1(RNAi)* animals in a normal regeneration assay (Figure S6A). We found that *zeb-1(RNAi)* also led to a failure to regenerate correctly in our migration assay (Figure S6B). Subsequent WFISH experiments revealed clear defects in cell migration and cell process formation, very similar to those observed for both *snail* TFs (Figure 7J-Q, Figure S6H and I). While we could only detect *zeb-1* transcript expression in relatively few migrating *smedwi-1^+^* NBs (8%, Figure S6C-F), this seems likely to be partly due to very low levels of transcript expression (Figure S6C-F) (Kao et al, 2017). Taken together, our data establish that conserved EMT-TFs are required for NB and progeny migration in planarians, establishing conservation of this regulatory circuit across bilaterians.

## DISCUSSION

### An X-ray shielded assay allows precise observation of cell migration and application of functional genomic approaches

We have established a robust and reliable method that allows the regenerative planarian model system to be used to study cell migration. During homeostasis as well as standard regeneration experiments, NBs and stem cell progeny are always close to where they are required. Nonetheless, as with all metazoans, NBs and progeny must still move into the correct functional positions in the tissues and organs. In the case of very anterior region and the pharynx of the planarian body plan, that are devoid of NB, homeostasis must be achieved by migration of stem cell progeny (Figure S2A and B). However precise monitoring of this process is difficult as the migratory distances involved are short and so confidently inferring changes in migratory behavior as oppose to changes in, say, differentiation is not possible. Our X-ray shielded assay creates a ‘blank canvas’ into which migrating stem cells and stem cell progeny move and we can accurately assign relationships between migration, differentiation and proliferation of groups of these cells over time. We show that planarian NBs and progeny are capable of restoring full tissue and organ function by migrating from the small shielded region. The innovations we have made here compared to earlier approaches allow for a thinner shield, smaller worms to be irradiated and technical consistency over relatively large numbers of worms. This has allowed us to combine WFISH and RNAi approaches so that we can now use the planarian model to study migration in a regenerative context.

### A detailed description of migratory behavior in a regenerative context

In this work we have revealed a number of detailed properties of cell migration in planarians that can be used to help unpick the mechanisms controlling cell migration. We have shown that migration occurs in response to wounding or damaged tissue as previously described (Guedelhoefer and Sánchez Alvarado, 2012a). We also find that contrary to previous work that migration can occur without wounding or failure in tissue homeostasis for anteriorly positioned stem cells and stem cell progeny. This observation tallies with the absence of NBs in anterior regions and the brain in intact animals, which suggests that a mechanism for encouraging homeostatic cell migration must exist. We also observe that migrating cells home precisely to wounds without initially recognizing other tissue regions also lack NBs and progeny. Finally, we observe that in regions containing moving cells we can see a clear increase in the number of cells with pronounced cell extensions. Migrating cells are unconnected to other migrating cells, and together these observations give an EMT like characteristic to planarians cell migration, as oppose to other mechanism involving collective cell migration. Taken together these observations establish a set of basic phenotypic criteria that can be used to the study the genetic and molecular control of cell migration.

### The relationship between stem cell migration, proliferation and differentiation

Stem cell migration during normal healthy tissue homeostasis must be intricately linked to cell divisions, differentiation and integration of new cells to ensure dysfunctional aged and damaged differentiated cells are successfully replaced. Studying this process *in vivo* during adult tissue homeostasis has proven to be challenging and remains limited to a few contexts. Highly regenerative animal models represent an opportunity to study these processes, which together power regeneration. Thus, perhaps the most important observations facilitated by our assay are those concerning the relationships between migration, proliferation and differentiation.

We observe that progeny migrate in large numbers in an initial response to wounding and that proliferating NBs accompany them in smaller numbers. In response to both wounding and homeostatic signals we observe that NBs divide asymmetrically as they migrate, and that the new progeny differentiate further while they migrate. For the well-characterised epidermal lineage this creates an order of migration that recapitulates the order of differentiation. We do not see any evidence that progeny slow down their differentiation process in order to first reach wound sites and then differentiate. Instead our observations broadly recapitulate cell behaviour observed during regeneration, in which progeny migrate to form the regeneration blastema where they complete differentiation and cycling cells follow later. Our analysis detects significant differences in migration between *smedwi-1^+^* cells and zeta class*/smedwi-1^-^* cells, which we interpret as suggesting that newly minted progeny migrate ahead of cycling NBs as they do in blastema formation. NBs may migrate more slowly on average as they stop to divide, or because they require the presence of progeny at certain density before they can be healthily maintained in a repopulating tissue region, or simply perhaps because they are slower due to having smaller cell extensions. Based on these observations we note that our assay will provide an alternate method of assessing cell lineage relationships with WFISH approaches and when combined with RNAi it allows the molecular processes controlling the interplay between migration, proliferation and differentiation to be studied. For example, future experiments can test the requirement of migrating stem cells for stem cell progeny by interfering with differentiation of specific lineages or asymmetric division.

Related to the observation that the order of cell migration we observe recapitulates cell lineages is the question of whether all types of wound will result in the same or different combinations of migratory, proliferative and differentiation responses. While we have established that migration homes precisely to wound sites we can also now study if differentiation programs show specificity to the type of wounds depending on which cell types are damaged. Recently it was shown that production of photoreceptor precursors and cells was independent of whether eyes were present or not (LoCascio et al., 2017), suggesting that for some these organs at least differentiation programs are independent of the state of the target tissue in planarians. Combining our assay with experimental paradigms that damage one or a few defined cell types will help begin to answer how demands for new cells are regulated and how stem cells and their progeny sense and adjust to these demands. Given that these are likely to be the processes that decline in human age related disease or are mis-regulated during tumour progression, new planarian experiments in this context will provide important insights.

### A role for *notum* in homeostatic migration of stem cells and stem cell progeny

The precise identification of the signals that trigger migration after wounding remains an open question. It seems more than likely that many overlapping signals cooperate to ensure migration occurs correctly and they may include signals associated primarily with occurrence of damage as well as signals from specific tissues that require specific progeny. Two genes that have already been shown to have complementary roles in controlling the polarity of planarian regeneration, *wnt1* and *notum*, are both known to be wound induced (Petersen and Reddien, 2011) and represented good candidates for potential roles in cell migration after wounding. In addition homeostatic expression of *notum* was recently shown to be involved in regulating planarian brain size in combination with *wnt11-6*, and specifically ensuring that sufficient neural precursors are produced to maintain the correct brain size (Hill and Petersen, 2015). These observations therefore also made *notum* a candidate for involvement in the homeostatic cell migration that we described in intact animals in anterior regions.

Using RNAi we found no role for either *wnt1* or *notum* in wound induced migration, however we found that *notum* is required for the homeostatic anterior migration. Given the homeostatic expression of *notum* transcript and the observation that cells migrate homeostatically within a certain distance from the anterior tip of the animal, we propose that a gradient of *notum* somehow provides directional cues to migrating cells. We note that, the formation of cell extensions is not effected by *notum(RNAi)*, suggesting that other signals may be responsible for this aspect of migratory behaviour while *notum* activity provides a directional cue. *Notum* in planarians, mammals and flies has been implicated as a Wnt signaling inhibitor (Kakugawa et al., 2015; Traister et al., 2008; Zhang et al., 2015), so it is possible that inhibition of local homeostatic levels of Wnt signaling, specifically of anteriorly expressed planarian Wnts (*wnt11-6* and *wnt5*) may then allow homeostatic migration. Future work with our assay will aim to understand the mechanism by which *notum* facilitates homeostatic migration and wound induced migration.

### Conservation of EMT-TF function and the potential to study processes relevant to tumor invasion

The fact that cells appear to migrate individually and that in migratory regions increased numbers of cells have extended cell processes suggested molecular mechanisms associated with EMT may regulate migration. In order to begin to test this possibility we investigated the function of two planarian *Snail* family transcription factors and a planarian ortholog of *zeb-1*, as these are conserved positive regulators of cell migration during EMT, required to down regulate the expression of genes that encode proteins that maintain cell-cell contacts, like E-cadherin (Thiery and Sleeman, 2006). Enhanced *snail* gene expression has reported in several different cancer types including ovarian carcinoma (Davidson et al., 2012), breast tumours (Blanco et al., 2002; Elloul et al., 2005); gastric cancers (Peng et al., 2014; Rosivatz et al., 2002); hepatocellular carcinomas (Miyoshi et al., 2005; Sugimachi et al., 2003); colon cancers (Pálmer et al., 2004) and synovial sarcomas (Saito et al., 2004). Overexpression or down regulation of *Snail* has shown to modulate invasiveness and metastasis in in vitro cancer cell culture studies (Adhikary et al., 2014; Belgiovine et al., 2016; Fan et al., 2012; Horvay et al., 2015; Sharili et al., 2013; Smith et al., 2014; Villarejo et al., 2015). Similarly, *zeb1* over-activity has also been implicated in tumorigenesis. These reports clearly suggest that EMT-TFs are key players in cancer invasion and metastasis.

Within the context of our assay RNAi of all three of these genes led to failure in cell migration and we were able to clearly observe decrease in cells showing extended cell processes, indicative of migratory morphology. Our data confirm the role of EMT-TFs in controlling migration in the context for our assay and suggest we can use this as a basis for studying EMT related processes in planarians. By combining functional approaches with expression screens starting with planarian homologs of EMT-related transcription factor regulators and known upstream EMT regulatory signals, we will be able to find out more about EMT in the context of tissue homeostasis, regeneration and adult stem cell activity.

## EXPERIMENTAL PROCEDURES

### Planarian culture

A *Schmidtea mediterranea* asexual strain was cultured and maintained in 0.5% instant ocean water in the dark at 20^o^C. Animals were starved at least 7 days before using for experiments.

### X-ray irradiation, and design of shield

Irradiations were performed using a Comet MXR-321 x-ray set operated at 225 kVp, 17mA with a 0.5 mm aluminium filter. The X-ray field is collimated to 40 mm x 20 mm with a 6.1 mm thick lead disc positioned centrally, directly above the X-ray tube focal spot and supported within an aluminium frame. The removable central shielded area is achieved using a 0.8 mm wide, 6.1 mm thick lead strip spanning the long axis of the collimated field, this sits slightly proud of the main lead collimator so that it is in contact with the base of the Petri dish. When in position, the worms are irradiated at a dose rate of 23 Gy/min, reducing to ∼ 1 Gy/min underneath the shielded region. The variation in dose distribution across the strip is shown in supplementary figure 1C. The circular hole in the top aluminium plate corresponds to the outside diameter of the Petri dish and enables dishes to be positioned quickly and reproducibly. Thin strips of materials such as tungsten or tantalum could be used to replace the lead strip to achieve thinner shielded regions if required.

### Dosimetry

Dose rate measurements and spatial characterization of the treatment field was performed using Gafchromic EBT3 film (International Specialty products, Wayne, NJ) placed in the base of an empty 60 mm Petri dish. Twenty-four hours following exposure the EBT3 film was scanned in transmission mode at 48 bit RGB (16 bits per colour) with 300 dpi resolution using a flatbed scanner (Epson Expression 10000XL). A template was used to position the film within the scanner and the scanning direction was kept constant with respect to the film orientation, as recommended in the manufacturer’s guidelines. The dose was calculated using the optical density of the red channel and corrected using the optical density of the blue channel in order to compensate for small non-uniformities in the film which cause false apparent variations in dose (as described in the technical brief: *Gafchromic EBT2 Self-developing film for radiotherapy dosimetry*). The batch of EBT3 film was calibrated following the recommendations of the report of AAPM Task Group 61(Ma et al., 2001).

### Shielded irradiation

Up to 20 size-matched planarians (3 to 4 mm) were anesthetized in ice cold 0.2% chloretone and aligned on 60mm Petri dish (Guedelhoefer and Sánchez Alvarado, 2012b). Anterior tip of all worms were aligned in a perfect line to keep the absolute migratory distance (distance between tip of the head and shielded region) fixed. Petri dish is pre-marked with a line at bottom denoting the place and dimensions (length and thickness) of the shield strip. Excess liquid is removed to minimize movement of worms during at the time of irradiation. Petri dish containing worms is then placed on to the shield of bottom source X-ray irradiator. Care is taken to perfectly match the position of shield trip and line marked on Petri dish to ascertain the exact region of the worm to be shielded. 30Gy X-ray (225kV for 1 min 18 seconds) is used for irradiation. Once irradiation is over, planarians were immediately washed with instant ocean water and transferred into fresh instant ocean water and incubated in dark at 20^o^C.

### WFISH, immunostaining and imaging

Whole mount fluorescent in-situ hybridization was performed as described previously (Currie et al., 2016; King and Newmark, 2013). H3ser10p rabbit monoclonal antibody from Millipore (04–817) was used for immunostaining (Felix and Aboobaker, 2010). Confocal imaging was done with Olympus FV1000 and Zeiss 880 Airyscan microscope. Bright field images were taken with Zeiss Discovery V8 from Carl Zeiss using Canon 1200D camera. Images were processed with Fiji and Adobe Photoshop. ZEN 2.1 (blue edition) software from Carl Zeiss was used to construct 3D images of cells. All measurements and quantifications were done with Fiji and Adobe Photoshop. Significance was determined by unpaired 2-tailed Student’s t-test.

### Gene cloning and RNAi

Planarian genes were cloned into the pPR-T4P plasmid vector containing opposable T7 RNA polymerase promoters (kind gift from Jochen Rink). The cloned vectors were then used for in vitro dsRNA synthesis and probe synthesis as described previously (King and Newmark, 2013; Rouhana et al., 2013). The primers used to generate dsRNA template from genes were as follows:

*mmpa* (GenBank: HE577120.1): Fw 5’-ATCCTGATTACGGCTCCAA-3’ and Re 5’-TTTATTGGGGGTGCAACTGT-3’

β*1-integrin* (GenBank: KU961518.1): Fw 5’-GAACTCAACACACAACGCCC-3’ and Re 5’-TCTCGACAGGGAACAATGGC-3’

*snail-1* (GenBank: XXXXX): Fw 5’-AGCAATCAATCCTAAAGTCG-3’ and Re 5’-CGATAGATTCTTCCACGGAG-3’

*snail-2* (GenBank: KJ934814.1): Fw 5’-GTTATCAAGCCAGACCTTCA-3’ and Re 5’-GTTTGACTTGTGAATGGGTC-3’

*zeb-1* (GenBank: XXXXX): Fw 5’-TCGTACCCTCATCTACCGCA-3’ and Re 5’-GGGTTTCTCTCCGCTGTGAA-3’

Previously described sequence regions were used for dsRNA synthesis of *wnt1* (Petersen and Reddien, 2009) and *notum* (Petersen and Reddien, 2011). Reported sequences were used for riboprobe synthesis of *smedwi-1* (Reddien et al., 2005), *prog-1* (Eisenhoffer et al., 2008), *agat-1* (Eisenhoffer et al., 2008), zeta pool (van Wolfswinkel et al., 2014), and sigma pool (van Wolfswinkel et al., 2014). To generate probes for *mmpa,* β*1-integrin, snail-1, snail-2* and *zeb-1* the same regions of their respective dsRNA were used. For knockdown of genes animals were injected with 3 x 32nl of dsRNA 6 times over 2 weeks. If worms need to be used for shielded irradiation after RNAi, a 1-day gap was kept between last RNAi injection and the shielded irradiation.

## SUPPLEMENTAL INFORMATION

Supplemental Information includes six supplemental figures.

## AUTHOR CONTRIBUTIONS

PA and AAA designed the experiments. PA, EA, NK performed the experiments. JT and MH helped with designing X-ray shield and performing X-ray shielded irradiations. PA and AAA wrote the manuscript.

## ACKNOWLEDGMENTS

We thank all members of the Aboobaker lab past and present for discussions and reagent sharing. The work of PA, EA, NK, AAA is funded by the MRC (grant number MR/M000133/1), BBSRC (grant number BB/K007564/1), the John Fell Fund Oxford University Press (OUP), and a small grant from the CRUK Oxford Centre. NK is funded by a Marie Curie Sklodowska fellowship funded by Horizon 2020. MH and JT acknowledge funding from the Funding from Medical Research Council Strategic Partnership Funding (MC-PC-12004) for the CRUK/MRC Oxford Institute for Radiation Oncology is gratefully acknowledged.

## SUPPLEMENTARY FIGURE LEGENDS

**Figure S1. Parts and dimensions of lead shield assembly**

(A) Lead strip and lead shield are assembled with aluminium support which further covered with aluminium disc to support Petri dish in the final lead shield assembly.

(B) Dimensions of lead shield and lead strip from top and side view. Unit: mm.

(C) Dose distribution across the lead strip showing greater than 95% attenuation of X-ray dose under the shield protected region.

**Figure S2. General features of cell migration and different shapes of migrating and non-migrating cells**

(A) FISH showing distribution of stem cells (green) in intact wild type worm. Stem cells are absent in the pharynx region, in brain region and region anterior to photoreceptors (*). Scale bar: 500μm.

(B) FISH showing that stem cells (green) are absent in the early regenerative blastema in a tail fragment regenerating at 3dpa (n=5). Scale bar: 200μm.

(C) H3P immunostaining shows increase in mitotic cells (yellow) in the migratory region in decapitated animals over the time course, 1dpa, 4dpa, 7dpa and 10dpa (n=5 per time point). Scale bar: 500μm.

(D) Graph showing increasing distance of mitotic cells (magenta dots) from the shielded region over the time course, 1dpa, 4dpa, 7dpa and 10dpa (n=5 per time point). Each dot represents the distance of individual H3P cell from the shielded region. 5 most distal H3P cells were considered for measurements from each animal. Lines and error bars indicate mean and SD.

(E, F) Stem cells (green) and early progeny (magenta) show directional migration towards the site of poking (E) and notch (F).

(G) Stem cells (green) and early progeny (magenta) from the dorsal side migrate more rapidly than the ventral side. Scale bar: 100μm.

(H) Measurements of distance migrated by stem cells (green) and early progeny (magenta) from dorsal and ventral side in decapitated animal at 4dpa. Each dot represents average the distance migrated by 10 most distal cells in an animal (n=5). Lines and error bars indicate mean and SD.

(I) Montage showing migrating stem cells (green) in different planes from dorsal to ventral side (1 to 6).

(J-K) Different morphology of stem cells (green) (J) and early progeny (magenta) (K) without and with extended processes.

**Figure S3. Regenerative morphology of RNAi animals and expression patterns of *mmpa* and β*1-integrin***

(A) Head, Trunk and Tail fragments regenerated at 11 days post amputation following *gfp(RNAi)*, *mmpa(RNAi)* and β*1-integrin(RNAi)*. (n=10)

(B) Rescue and regeneration of *gfp(RNAi)*, *mmpa(RNAi)* and β*1-integrin(RNAi)* worms following shielded irradiation and decapitation. (n=30)

(C-D) Expression (C) and FPKM (D) profile of *mmpa* in X1, X2 and Xins cell population.

(E) FISH showing whole body expression pattern of *mmpa*.

(F-G) FISH showing expression of *mmpa* in *smedwi-1^+^* NBs (F) and *prog-1^+^* progeny (G) at 2dpa. Around 3% *smedwi-1^+^* NBs express *mmpa* and no detectable expression of *mmpa* found in *prog-1^+^* progeny. Scale bars: 20μm

(H-I) Expression (H) and FPKM (I) profile of β*1-integrin* in X1, X2 and Xins cell population.

(J) FISH showing whole body expression pattern of β*1-integrin*.

(K-L) FISH showing expression of β*1-integrin* in *smedwi-1^+^* NBs (K) and *prog-1^+^* progeny (L) at 2dpa. Around 92% *smedwi-1^+^* NBs and 33% *prog-1^+^* progeny express β*1-integrin*.

(M) FISH shows stem cells (green) and early progeny (magenta) migrate and repopulate the entire migratory region at 15dpa in *gfp(RNAi)* animals but the migration is inhibited in *mmpa(RNAi)* and β*1-integrin(RNAi)* worms that leads to regression of anterior tissue.

(N) Measurements shows drastic decrease in the distance migrated by stem cells (green) and early progeny (magenta) at 15dpa in *mmpa(RNAi)* and β*1-integrin(RNAi)* animals compared to *gfp(RNAi)* worms (n=5). Each dot represents the average distance migrated by 10 most distal cells from each animal. Lines and error bars indicate mean and SD. Student’s t test: *p<0.05.

**Figure S4. Regenerative phenotype of *notum* and *wnt1* RNAi animals**

(A) Head, Trunk and Tail fragments regenerated at 11 days post amputation following *gfp(RNAi)*, *notum(RNAi)* and *wnt1(RNAi)*. (n=10)

(B) Rescue and regeneration of *gfp(RNAi)*, *notum(RNAi)* and *wnt1(RNAi)* worms following shielded irradiation and decapitation. (n=30)

(C) Rescue of intact uninjured animals in *gfp(RNAi)* and *notum(RNAi)* worms following shielded irradiation. (n=30)

(D) FISH shows stem cells (green) and early progeny (magenta) migrate anteriorly and repopulate almost entire migratory region at 20dpi in *gfp(RNAi)* animals but the migration is inhibited in *notum(RNAi)* worms that leads to regression of anterior tissue.

(E) Measurements shows drastic decrease in the distance migrated by stem cells (green) and early progeny (magenta) at 20dpi in *notum(RNAi)* compared to *gfp(RNAi)* worms (n=5). Each dot represents the average distance migrated by 10 most distal cells from each animal. Lines and error bars indicate mean and SD. Student’s t test: *p<0.05.

**Figure S5. Regenerative morphology of RNAi animals and expression patterns of *snail-1* and *snail-2***

(A) Head, Trunk and Tail fragments regenerated at 11 days post amputation following *gfp(RNAi)*, *snail-1(RNAi)* and *snail-2(RNAi)*. (n=10)

(B) Rescue and regeneration of *gfp(RNAi)*, *snail-1(RNAi)* and *snail-2(RNAi)* worms following shielded irradiation and decapitation. (n=30)

(C-D) Expression (C) and FPKM (D) profile of *snail-1* in X1, X2 and Xins cell population.

(E) FISH showing whole body expression pattern of *snail-1*.

(F-G) FISH showing expression of *mmpa* in *smedwi-1^+^* NBs (F) and *prog-1^+^* progeny (G) at 2dpa. Around 87% *smedwi-1^+^* NBs express *snail-1* and very little (∼1%) expression of *snail-1* found in *prog-1^+^* progeny. Scale bars: 20μm.

(H-I) Expression (H) and FPKM (I) profile of *snail-2* in X1, X2 and Xins cell population.

(J) FISH showing whole body expression pattern of *snail-2*.

(K-L) FISH showing expression of *snail-2* in *smedwi-1^+^* NBs (K) and *prog-1^+^* progeny (L) at 2dpa. Around 93% *smedwi-1^+^* NBs and less than 3% *prog-1^+^* progeny express *snail-2*. Scale bars: 20μm.

(M) FISH shows stem cells (green) and early progeny (magenta) migrate and repopulate the entire migratory region at 15dpa in *gfp(RNAi)* animals but the migration is inhibited in *snail-1(RNAi)* and *snail-2(RNAi)* worms that leads to regression of anterior tissue.

(N) Measurements shows drastic decrease in the distance migrated by stem cells (green) and early progeny (magenta) at 15dpa in *snail-1(RNAi)* and *snail-2(RNAi)* animals compared to *gfp(RNAi)* worms (n=5). Each dot represents the average distance migrated by 10 most distal cells from each animal. Lines and error bars indicate mean and SD. Student’s t test: *p<0.05.

**Figure S6. Effect of *zeb-1* RNAi on regeneration and it’s expression in different cell population**

(A) Head, Trunk and Tail fragments regenerated at 11 days post amputation following *gfp(RNAi)* and *zeb-1(RNAi)* animals. (n=10)

(B) Rescue and regeneration of *gfp(RNAi)* and *zeb-1(RNAi)* worms following shielded irradiation and decapitation. (n=30)

(C-D) Expression (C) and FPKM (D) profile of *zeb-1* in X1, X2 and Xins cell population.

(E) FISH showing whole body expression pattern of *zeb-1*.

(F-G) FISH showing expression of *zeb-1* in *smedwi-1^+^* NBs (F) and *prog-1^+^* progeny (G) at 2dpa. Around 8% *smedwi-1^+^* NBs express *zeb-1* and very little (∼1%) expression of *zeb-1* found in *prog-1^+^* progeny. Scale bars: 20μm.

(H) FISH shows stem cells (green) and early progeny (magenta) migrate and repopulate the entire migratory region at 15dpa in *gfp(RNAi)* animals but the migration is inhibited in *zeb-1(RNAi)* worms that leads to regression of anterior tissue.

(I) Measurements shows drastic decrease in the distance migrated by stem cells (green) and early progeny (magenta) at 15dpa in *zeb-1(RNAi)* animals compared to *gfp(RNAi)* worms (n=5). Each dot represents the average distance migrated by 10 most distal cells from each animal. Lines and error bars indicate mean and SD. Student’s t test: *p<0.05.

